# Human primary astrocytes increase basal fatty acid oxidation following recurrent low glucose to maintain intracellular nucleotide levels

**DOI:** 10.1101/271981

**Authors:** Paul G Weightman Potter, Julia M. Vlachaki Walker, Josephine L Robb, John K. Chilton, Ritchie Williamson, Andrew Randall, Kate L.J. Ellacott, Craig Beall

**Affiliations:** Institute of Biomedical and Clinical Sciences, University of Exeter Medical School, RILD Building, Barrack Road, EX2 5DW, UK; School of Pharmacy and Medical Sciences, University of Bradford, BD7 1DP, UK; Hatherly Laboratories, Prince of Wales Road, University of Exeter, EX4 4PS, UK

## Abstract

Hypoglycemia is a major barrier to good glucose control in type 1 diabetes and frequent exposure to hypoglycemia can impair awareness to subsequent bouts of hypoglycemia. The neural changes that occur to reduce a person’s awareness of hypoglycemia are poorly defined. Moreover, the molecular mechanisms by which glial cells contribute to hypoglycemia sensing and glucose counterregulation require further investigation. To test whether glia, specifically astrocytes, could detect changes in glucose, we utilized human primary astrocytes (HPA) and U373 astrocytoma cells and exposed them to recurrent low glucose (RLG) *in vitro*. This allowed measurement, with high specificity and sensitivity, of changes in cellular metabolism following RLG. We report that the AMP-activated protein kinase (AMPK) is activated over a pathophysiologically-relevant glucose concentration range. We observed an increased dependency on fatty acid oxidation for basal mitochondrial metabolism and hallmarks of mitochondrial stress including increased proton leak and reduced coupling efficiency. Relative to glucose availability, lactate release increased during low glucose but this was not modified by RLG, nor were glucose uptake or glycogen levels. Taken together, these data indicate that astrocyte mitochondria are dysfunctional following recurrent low glucose exposure, which could have implications for hypoglycemia glucose counterregulation and/or hypoglycemia awareness.

## INTRODUCTION

Hypoglycemia in type 1 diabetes and advanced insulin-treated type 2 diabetes remains a significant concern for people with diabetes. For patients, it is a major barrier to enjoying the health benefits of exercise [1, 2] and is increasingly thought to contribute directly to an increased risk of cardiovascular events in type 1 and type 2 diabetes [3, 4]. Individuals that experience frequent hypoglycemia or have had long-standing diabetes, produce a diminished counterregulatory response to hypoglycemia [5, 6], often accompanied by reduced symptom awareness, termed “impaired hypoglycemia awareness”.

Defective glucose counterregulatory responses to hypoglycemia is, at least in part, mediated by changes in the function of critically important glucose-sensing (GS) neurons in the ventromedial nucleus of the hypothalamus (VMH) playing an important role; although GS neurons in the hindbrain have also been implicated [7]. In addition to neurons, the brain contains at least as many cells that lack electrically excitable membranes. Of these the largest group is the glial cells, the largest sub-population of which are astrocytes. As well as exhibiting a range of well-developed cell-to-cell signaling pathways, astrocytes are increasingly recognized as important players in central nervous system based-diseases such as obesity [8, 9], Alzheimer’s disease [10], Parkinson’s disease [11, 12] and motor neuron disease [13].

Recent studies have demonstrated that glucoprivic stimuli such as 2-deoxyglucose (2-DG) increase intracellular calcium in hindbrain astrocytes [14], which is a well-accepted marker of increased astrocyte activity. In the hypothalamus, astrocyte activation by fasting is indicated by increased glial fibrillary acidic protein (GFAP) expression and process complexity [15, 16], both markers of gliosis.

Astrocytes contain small amounts of glycogen that can be broken down to glucose and released as lactate to feed active neurons. For example, astrocytic glycogen stores can maintain axonal activity during energetic stress in white matter [17, 18] and also contribute to memory formation [19, 20]. Brain glycogen concentrations are increased on recovery from exercise [21], suggesting that astrocytic glycogen stores are both physiologically and pathophysiologically regulated. Astrocytes are in intimate apposition to the cerebral vasculature meaning that a change in glycemia may be detected by astrocytes before neurons. Moreover, astrocytes are generally accepted as metabolically flexible neural cells that can rapidly upregulate glycolysis to support neuronal function, partly due to the higher expression of 6-phosphofructo-2-kinase type 3(PFKFB3) that can stimulate glycolysis [22]. Despite this, little is known about the intrinsic response of human primary astrocytes to acute and recurrent low glucose (RLG) that occurs in type 1 diabetes. Given that astrocytes are metabolically flexible cells, we hypothesized that human astrocytes would 1) sense low glucose by increasing the phosphorylation of the cellular energy sensor AMP-activated protein kinase (AMPK) and 2) exhibit altered glycolytic and mitochondrial metabolism following RLG. To do this we utilized both human primary astrocytes and U373 astrocytoma cells and measured directly components of glycolytic and mitochondrial metabolism, lactate release, glycogen levels, intracellular nucleotide levels, and expression of key metabolic enzymes. We demonstrate for the first time that human astrocytes metabolically adapt by increasing basal fatty acid oxidation following a recurrent low glucose challenge that allows intracellular nucleotide levels to be maintained.

## RESEARCH DESIGN AND METHODS

### Astrocyte isolation and cell culture

Human primary astrocytes (HPA) were isolated from normal subventricular deep white matter blocks immediately post-mortem following consent from next-of-kin and with ethical approval from the North and East Devon Research Ethics Committee. Astrocytes were isolated from tissue blocks as previously described [23], sub-cultured and frozen stocks prepared. HPA cells were maintained in humidified incubators with 95% O_2_/5% CO_2_ in DMEM (ThermoFisher, #31885) containing 5.5 mmol/l glucose, supplemented with 10% FBS (Gibco, 10270-106) and gentamicin (25 μg/ml; ThermoFisher, 15750-060). U373 cells were maintained in DMEM (Sigma-Aldrich #D5671) containing 25 mmol/l glucose, supplemented with FBS (Gibco, 10270-106), and, penicillin and streptomycin (Gibco, 100 U/ml; 100 μg/ml). To enhance adherence of cells during growth, flasks and petri dishes containing HPA cells were pre-coated with poly-L-lysine (20 μg/ml; Sigma, P1274-25MG) in dH_2_O. 1 × 10^6^ cells were seeded into T75cm^2^ flasks and allowed to grow, and passaged after approximately 7-10 days. For experiments, cells were seeded in to 60 mm round petri dishes in media containing 5.5 mmol/l glucose with FBS, the day prior to study. On the morning of study, the glucose concentration was further reduced to 2.5 mmol/l glucose for 2 hours. Glucose concentrations were then dropped further to 0.1 mmol/l glucose as a hypoglycemic stimulus or maintained at 2.5 mmol/l glucose. For RLG studies over 4 days, cells were recovered in media containing high (5-5.5 mmol/l) glucose and FBS overnight (see Fig. S1 for culture model). The reasons for this were two fold; 1) higher glucose levels overnight (approximately 19 hours) prevented glucose depletion from the media, which would occur if the cells were maintained at 2.5 mM and 2) this pattern more accurately represented a model of glucose variation seen in type 1 diabetes.

### Immunoblotting

Cellular protein was isolated as previously described [24] with minor modifications. Briefly, for acute studies 450,000 cells, or for recurrent studies 250,000 cells were seeded on to 60 mm petri dishes and harvested at the end of experiments in 65 μl of lysis buffer. Protein concentrations were assessed by Bradford assay. 10 μg of protein was loaded per lane on 7-10% poly-acrylamide gels. Following electrophoresis, proteins were transferred to nitrocellulose membranes, blocked with BSA (5% w/v) or powered milk (5% w/v) and probed with antibodies against target proteins. Primary antibodies used were: pThr172 AMPK (1:1,000; #2535), pSer79 acetyl-CoA carboxylase (ACC; 1:1000; #3661), mouse α-actin (1:10,000; catalogue #A2066) was from Sigma Aldrich. Proteins were visualized and quantified using the Odyssey scanner (Licor, UK).

### Analysis of cellular metabolism

HPA and U373 cells were seeded at 2 × 10^5^ cells per well in Agilent Seahorse XF^e^96 assay plates the day prior to study. Media was exchanged for low-buffered media, pH manually adjusted to 7.40 at 37.0°C and cells placed in an atmospheric-CO_2_ incubator, to remove CO_2_ buffering capacity, one hour prior to starting the assay. Assay plates were loaded into the extracellular flux analyzer and studied on a 3-minute mix, 3-minute measure cycle and compounds injected as indicated. Following assay completion, the assay plate was removed from the analyzer, media removed and replaced with 100 μl of 50 mmol/l NaOH to lyse cells. Protein concentrations were determined by Bradford assay using NaOH as a vehicle. Cellular oxygen consumption (OCR) and extracellular acidification rates (ECAR) were normalized to total protein concentration of each well and where appropriate, normalized to baseline values (prior to compound injection). Mitochondrial stress tests (Agilent #103015-100) and mitochondrial fuel flexibility tests (#103270-100) were performed as per manufacturer’s instructions.

### Measurement of intracellular nucleotides

Total ATP levels were measured using ATPlite (Perkin Elmer; #6016941) with minor modifications as previously described [24]. ATP/ADP ratios were assayed by luminescence assay (Sigma; #MAK135). Briefly, 1×10^3^ cells were seeded on to black walled 96 well plates and exposed to 2.5 mmol/l or 0.1 mmol/l glucose levels for 15-180 minutes and nucleotide ratios assayed as per manufacturer’s instructions.

### Measurement of extracellular lactate, glucose uptake and glycogen levels

Extracellular lactate was measured by assessing NADH production from NAD^+^ in the presence of lactate dehydrogenase as previously described [25]. Briefly, 100 μL of extracellular supernatant was examined against a standard curve of lactate from 0-50 nM. Glucose uptake was measured using fluorescently labeled glucose analogue 6- (N-(7-Nitrobenz-2-oxa-1,3-diazol-4-yl)amino-6-Deoxyglucose (6-NBDG). 6-NBDG (600 μM) was added for 15 minutes in 2.5 mmol/l glucose in control conditions and in cells exposed to RLG. Glycogen levels were measured using a fluorometric kit from Cambridge Bioscience (Cambridge, UK) as per manufacturer’s instructions.

### Statistical analysis

For immunoblotting, densitometric values were normalized to unity to examine relative fold change in expression. A one-sample t-test was used to determine significant changes in phosphorylation or expression, relative to control. For comparisons of two groups a two-tailed unpaired t-test was used and for multiple group comparisons, a one-way ANOVA with *post-hoc* Bonferroni were used. To compare the mean differences between groups split by two independent variables, a two-way ANOVA with Bonferroni multiple comparisons tests was used. Statistical tests performed using GraphPad Prism software (Prism 5; GraphPad Software, La Jolla, CA, USA). Results are expressed as mean ± standard error, unless otherwise stated. *p* values of <0.05 were considered statistically significant.

## RESULTS

### AMP-activated protein kinase is activated by acute low glucose in human astrocytes

AMPK activation in neurons is required for sensing hypoglycemia [26], although it is not known whether astrocytic AMPK reacts to the same energy stress, therefore we exposed human astrocytes to normal (2.5 mmol/l) and low brain (0.1 mmol/l) glucose levels. In cells exposed to low glucose for 30 minutes, we noted increased phosphorylation of AMPK at threonine-172, a site required to be phosphorylated for full kinase activation in both the human primary astrocytes (HPA; Fig.1*A,C*) and U373 cells (Fig.1*B,D*). This was accompanied by increased phosphorylation of acetyl CoA carboxylase (ACC) at serine 79 (Fig.1*E,F*). These data are consistent with the ubiquitous role for AMPK in detecting low glucose levels in other cells types and now reported in glia.

**Figure 1.**
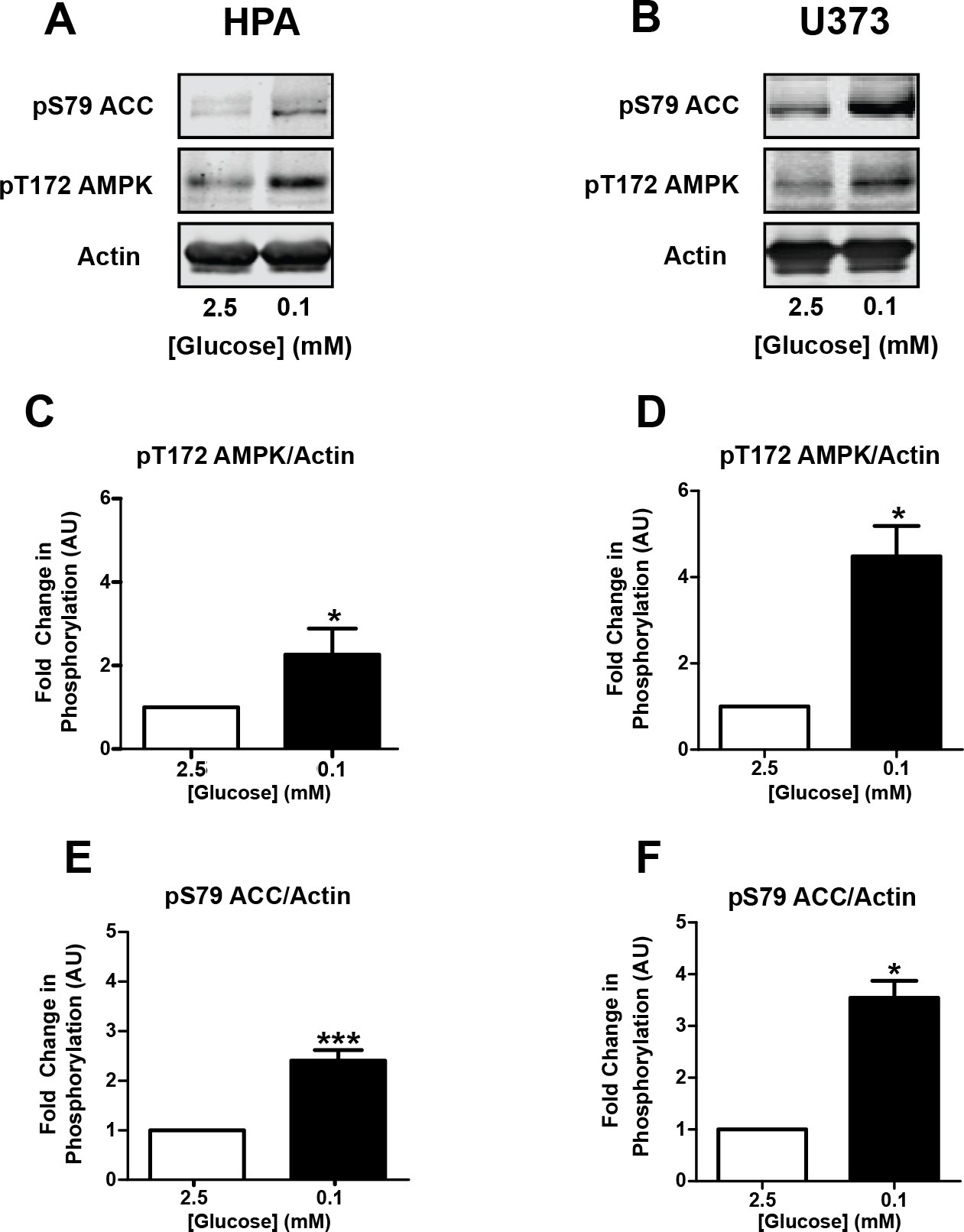
Acute low glucose increases AMP-activated protein kinase (AMPK) pathway activation. Representative immunoblots from human primary astrocyte (HPA, **A**; n=8) and U373 astrocytoma cells (**B**; n =3) exposed to 2.5 or 0.1 mmol/l glucose media for 30 minutes. ***C, D***. Densitometric analysis of immunostaining for pT172 AMPK normalized to actin as a loading control. ***E, F***. Densitometric analysis of immunostaining for pS79 ACC normalized to actin as a loading control. \**P*<0.05;\*\*\**P*<0.001. One-sample t-test in comparison to control.

### RLG exposure increases mitochondrial metabolism in human astrocytes

To determine whether baseline metabolism was altered following RLG, we performed mitochondrial stress tests on HPA and U373 cells following RLG (see Fig. S1 for RLG model). This was performed at euglycemic glucose levels post RLG to observe any cellular adaptations. Firstly, we observed significantly higher baseline oxygen consumption rate (OCR) in both HPA (Fig. 2*A,C*) and U373 cells (Fig. 2*B,D*) exposed to RLG (3 versus 0 prior bouts of low glucose), indicating persistent adaptations in cellular metabolism. Oligomycin was then added to block ATP synthase, FCCP to stimulate maximal respiration, and rotenone and antimycin A were added to inhibit complex I and III, respectively. This allowed calculation of coupling efficiency, which was significantly decreased in HPA cells only (Fig. 2*E,F*). However, both cell types displayed increased proton leak (Fig. 2*G,H*), indicating mitochondrial dysfunction. Despite the functional evidence of mitochondrial dysfunction, we saw no changes in the morphology of the filamentous mitochondrial networks between the treatment groups as examined using MitoTrackerRed (Fig. S2*A,B*).

**Figure 2.**
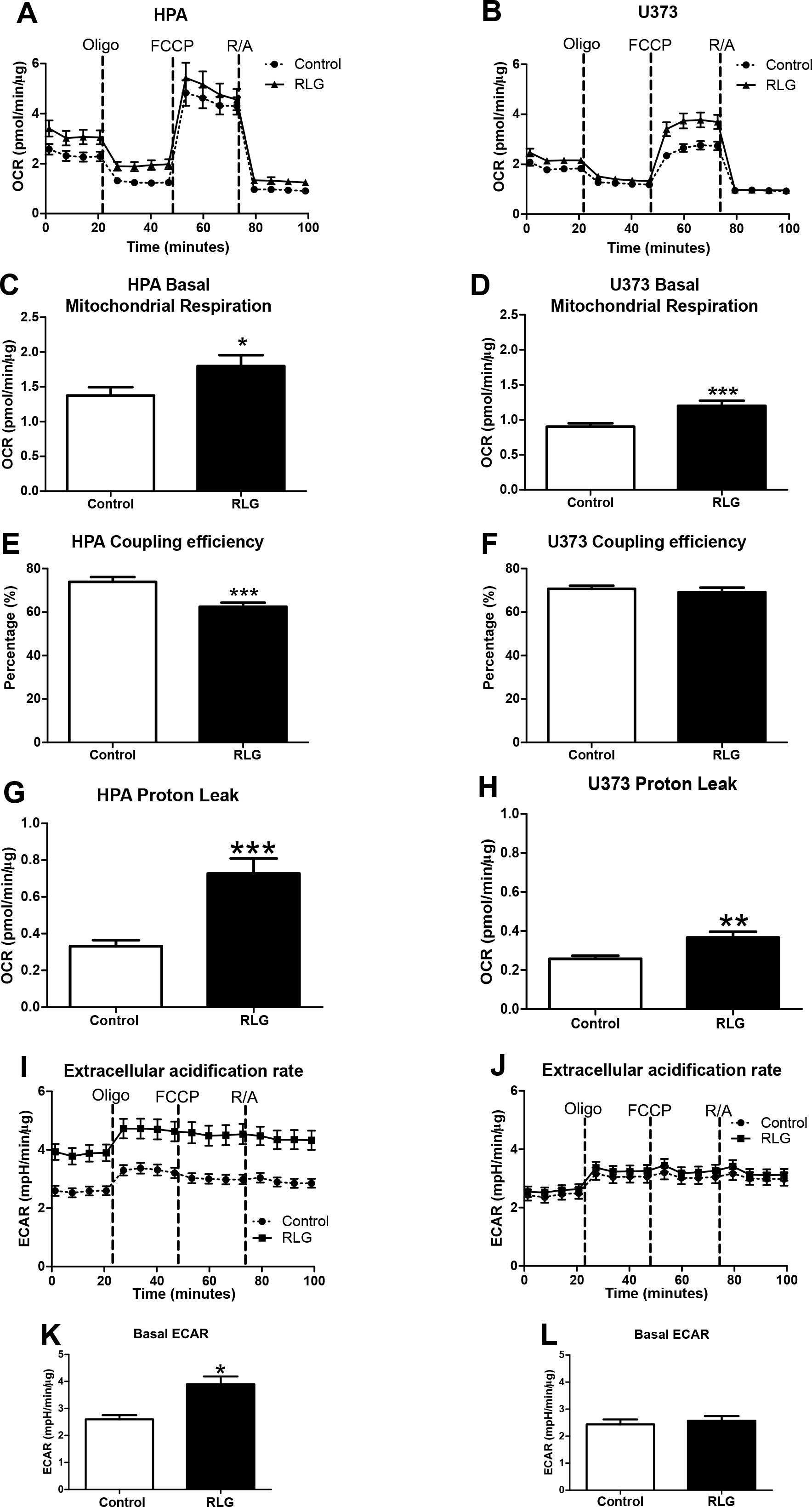
Recurrent low glucose (RLG) increases baseline mitochondrial metabolism and reduces mitochondrial coupling efficiency. Oxygen consumption rate (OCR) of human primary astrocytes (HPA; **A**; 36-39 across three separate assays) and U373 cells (**B**) following RLG (n=40-42, across across separate assays). Cells were exposed to oligomycin (10 μM), FCCP (5 μM) and a combination of rotenone and antimycin A (5 μM). ***C, D***. Mean basal OCR in control and RLG treated astrocytes. ***E, F***. Coupling efficiency calculated from the ratio of oligomycin sensitive OCR and basal OCR expressed as a percentage. ***G, H***. Proton leak, calculated from the oligomycin-insensitive (i.e. not ATP-synthase linked) OCR minus non-mitochondrial respiration, from HPA cells and U373 cells. ***I, J***. Extracellular acidification rate (ECAR) analysis measured during mitochondrial stress tests. ***K, L***. Mean baseline ECAR in HPA and U373 cells following RLG, respectively. \**P*<0.05;\*\*\**P*<0.001. Two-tailed students t-test.

In addition to changes in OCR, we also noted significantly elevated basal extracellular acidification rate (ECAR) in HPA cells exposed to RLG (Fig. 2*I,K*), indicating enhanced glycolysis following RLG. This did not occur within U373 cells following RLG (Fig. 2*J,L*).

### Astrocytes exposed to RLG display enhanced glycolysis on recovery of glucose level

Given the increased basal ECAR in HPA cells exposed to RLG, we next examined glycolytic re-activation after an acute glucose withdrawal (~80 minutes) in HPA cells following antecedent RLG. Under control conditions, we noted that the ECAR (glycolysis) response to glucose was concentration dependent (Fig. 3*A-C*), although ECAR was not significantly higher at 5.5 mmol/l glucose than 2.5 mmol/l glucose. Interestingly, the ECAR response to glucose in cells exposed to RLG (3 bouts of low glucose) was significantly greater than control (0 bouts of low glucose), indicating an enhanced glycolytic capacity (Fig. 3*A-C*). This only occurred at euglycemic and hyperglycemic levels. Consistent with enhanced glycolytic activity, we also noted a significant reduction in oxygen consumption rate (OCR) following glucose injection, indicative of a shift away from mitochondrial metabolism towards glycolysis (Fig.3*D-F*). Correspondingly, the reduction in OCR following glucose recovery was augmented in HPA cells exposed to RLG. Taken together these data suggest an enhanced Warburg effect in HPA cells following recovery of glucose levels. In U373 cells, RLG did not alter ECAR between groups (control vs RLG) on glucose recovery. However, the re-activation of ECAR on delivery of glucose was not concentration-dependent and in U373 cells the relative increase in ECAR was twice as large as that in HPA cells (Fig. S3). These studies show, obtaining primary cells from source is preferably to transformed cells.

**Figure 3.**
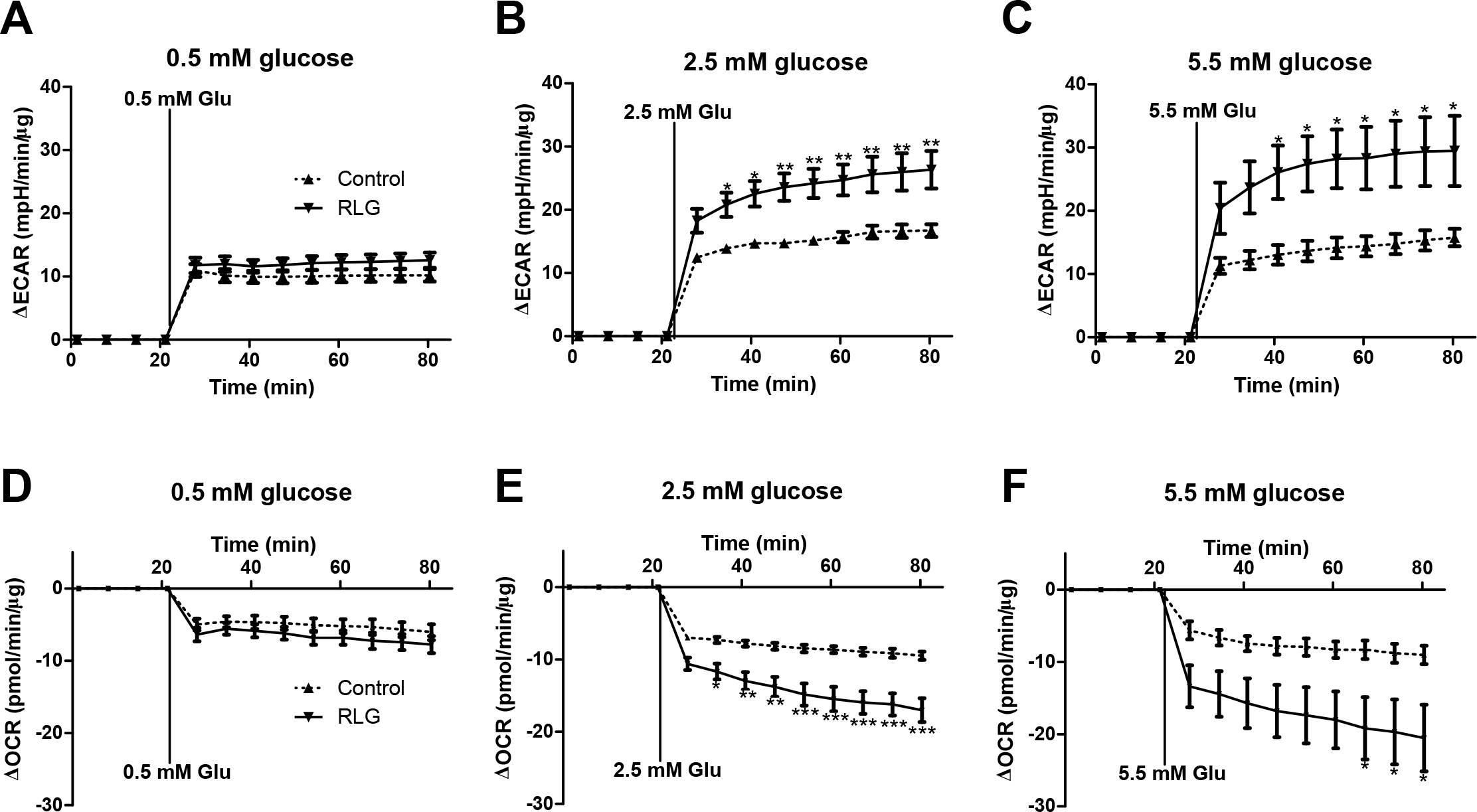
Recurrent low glucose (RLG) increases glycolytic re-activation after acute low glucose exposure. Extracellular acidification rate (ECAR) increased in human primary astrocyte (HPA) cells on re-introduction of 0.5 mmol/l glucose (**A**), 2.5 mmol/l glucose (**B**) and 5.5 mmol/l glucose (**C**). Oxygen consumption rate (OCR) decreased on addition of 0.5 mmol/l glucose (**D**), 2.5 mmol/l glucose (**E**) and 5.5 mmol/l glucose (**F**). \**P*<0.05; \*\**P*<0.01; \*\*\**P*<0.001 (n=8-10).

### Fatty acid oxidation (FAO) is increased in human astrocytes following RLG

To assess potential changes to mitochondrial fuel usage following RLG we performed mitochondrial fuel flexibility tests; achieved by sequential selective inhibition of the mitochondrial pyruvate carrier (MPC; glucose oxidation pathway), glutaminase (glutamine oxidation pathway) and carnitine palmitoyltransferase 1 (CPT1; fatty acid oxidation pathway). Surprisingly, there was no change in mitochondrial dependency, capacity or flexibility to metabolize pyruvate (Fig. 4*A-C*). This suggests that in astrocytes alterations to glucose metabolism are upstream of the transport of pyruvate into the mitochondria in astrocytes. There was a non-significant trend towards reduced glutamine dependency at baseline following RLG (Fig. 4*D*), with no significant change to the glutamine oxidation capacity (Fig. 4*E*). Combined, this produced a significant increase in glutamine flexibility (Fig. 4*F*). Interestingly, RLG significantly increased metabolic dependency on fatty acids (FA) but with no change in FAO maximum capacity, thereby causing a decrease in flexibility (Fig. 4*G-I*). These data demonstrate that RLG induces mitochondrial adaptations to fuel sources consistent with a shift towards increased reliance on FAO and an enhanced flexibility to utilize glutamine.

**Figure 4.**
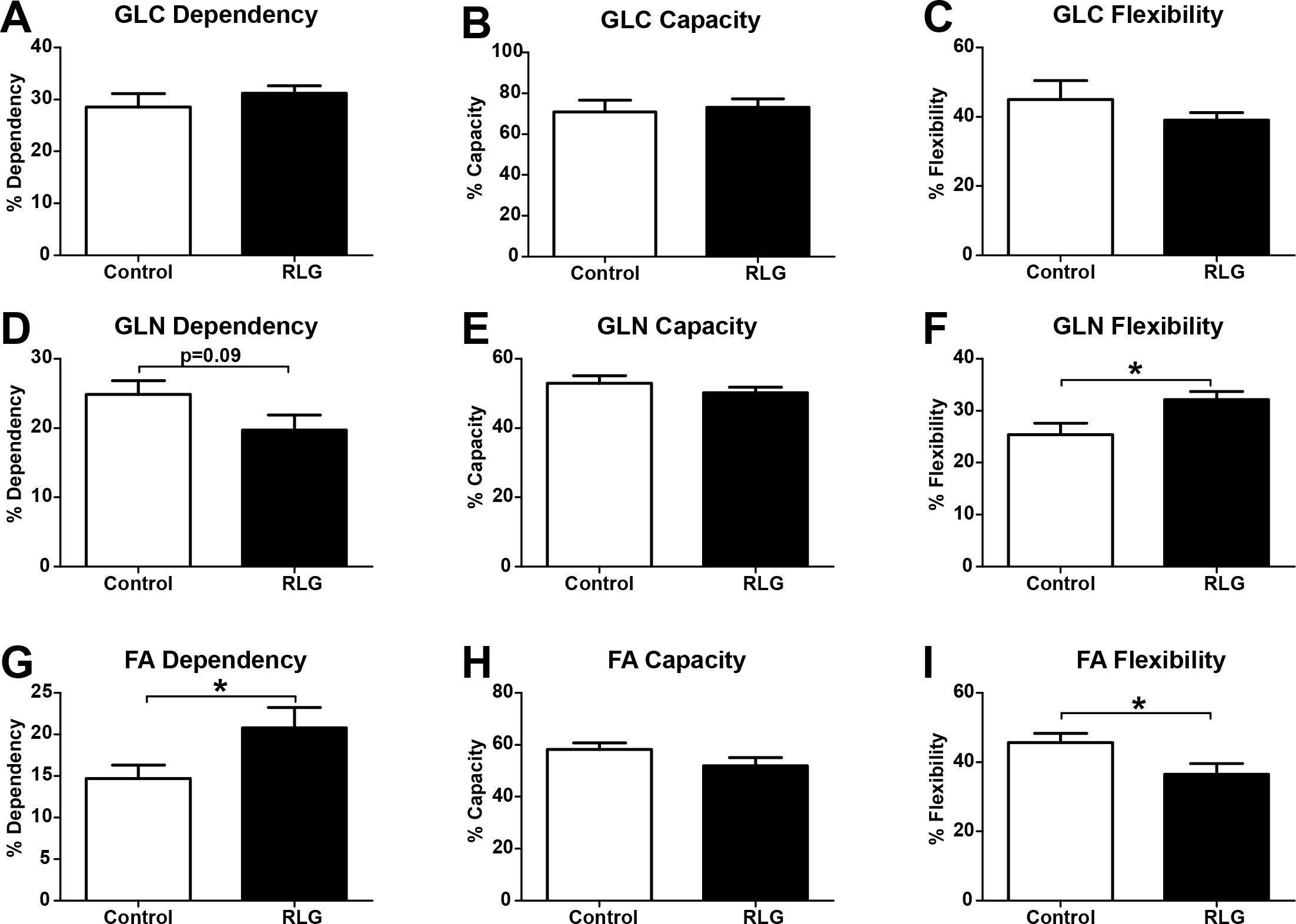
Recurrent low glucose (RLG) alters human primary astrocyte (HPA) mitochondrial fuel flexibility to increase dependency on fatty acids for basal oxidation. ***A-C***. Inhibition of mitochondrial pyruvate carrier using UK5099 (2 μM) to measure dependency, maximum capacity and flexibility within the glucose oxidation pathway. ***D-F***. Inhibition of the glutaminase using BPTES (3 μM) to measure changes in the glutamine oxidation pathway following RLG. ***G-I***. Inhibition of carnitine palmitoyltransferase 1 CPT1 using etomoxir (4 μM) to measure changes in the fatty acid (FA) oxidation pathway following RLG. \**P*<0.05 (n=21-32). Dependency: contribution of that pathway to baseline oxygen consumption rate. Capacity: maximum ability to oxidize through that pathway when two other oxidative pathways are inhibited. Flexibility: calculated from the difference between dependency and capacity.

### Human astrocytes maintain energy production (total ATP and ATP/ADP ratio) during low glucose conditions

To examine the consequences of acute low glucose and RLG for astrocyte energy content, we measured total ATP and the ATP/ADP ratio. These measurements were made both in the presence of acute low glucose and following RLG. Interestingly, we found that total ATP levels did not significantly change in HPA (Fig. 5*A*) or U373 cells (Fig. 5*B*) during acute low glucose exposure (0.1 mmol/l) lasting 3 hours. Despite the increased basal mitochondrial and glycolytic metabolism described above, total ATP levels following RLG were also comparable to control (Fig. 5*A,B*). Similarly, the ATP/ADP ratio was not significantly different following 3 hours of 0.1 mmol/l glucose, nor was the basal ratio altered following RLG (Fig. 5*C,D*). These data indicate that the total ATP content and ATP/ADP ratio are well defended in astrocytes, at least in response to up to 3 hours of low glucose, and that changes to glycolytic and mitochondrial metabolism may be compensatory in order to sustain intracellular nucleotide levels.

**Figure 5.**
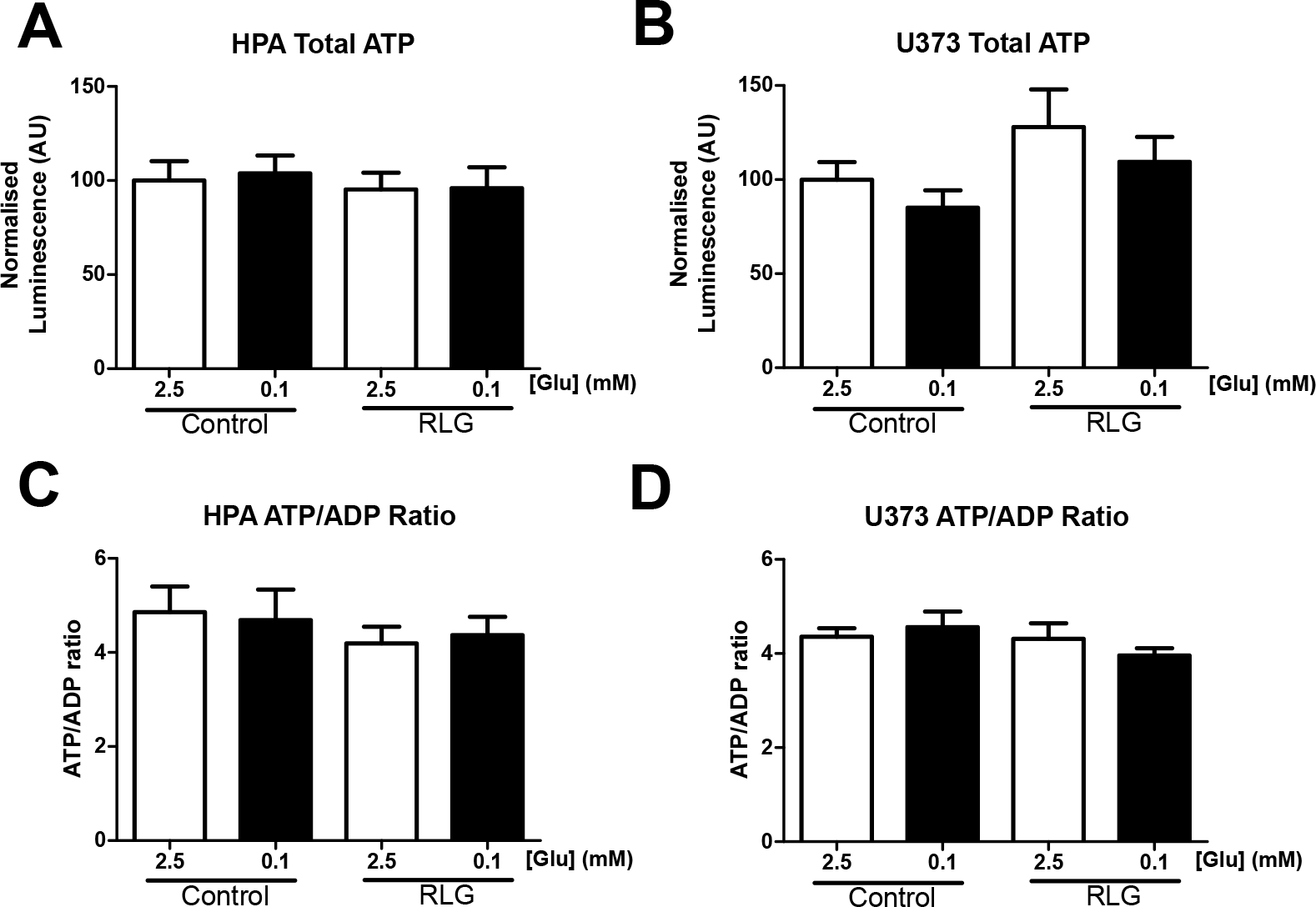
Acute and recurrent low glucose (RLG) does not modify intracellular ATP and ATP/ADP ratios. Total intracellular ATP levels in human primary astrocytes (HPA; **A**, n=6) and U373 astrocytoma cells (**B**; n=6) exposed to different glucose concentrations for 3 hours. Intracellular ATP/ADP ratios in HPA (**C**; n=6) and U373 astrocytoma cells (**D**; n=4) exposed to 2.5 or 0.1 mmol/l glucose for 3 hours.

### RLG does not alter glucose uptake or glycogen levels in human astrocytes

Western blotting demonstrated that the levels of key glycolytic markers GLUT1, hexokinase I (HKI), HKII, HKIII and phosphofructokinase (PFK) were not altered by acute or recurrent low glucose in human astrocytes (Fig. S4).

Lactate release from HPA and U373 cells decreased by 25-50% during 3 hours of low glucose exposure (Fig. 6*A,B*), although this was a substantial decrease, this was proportionally much less than the 25-fold reduction in glucose availability. Hence, when corrected for glucose availability, there was a large relative increase in lactate release (Fig. 6*C,D*); however, this was not different between control and RLG-treated cells.

**Figure 6.**
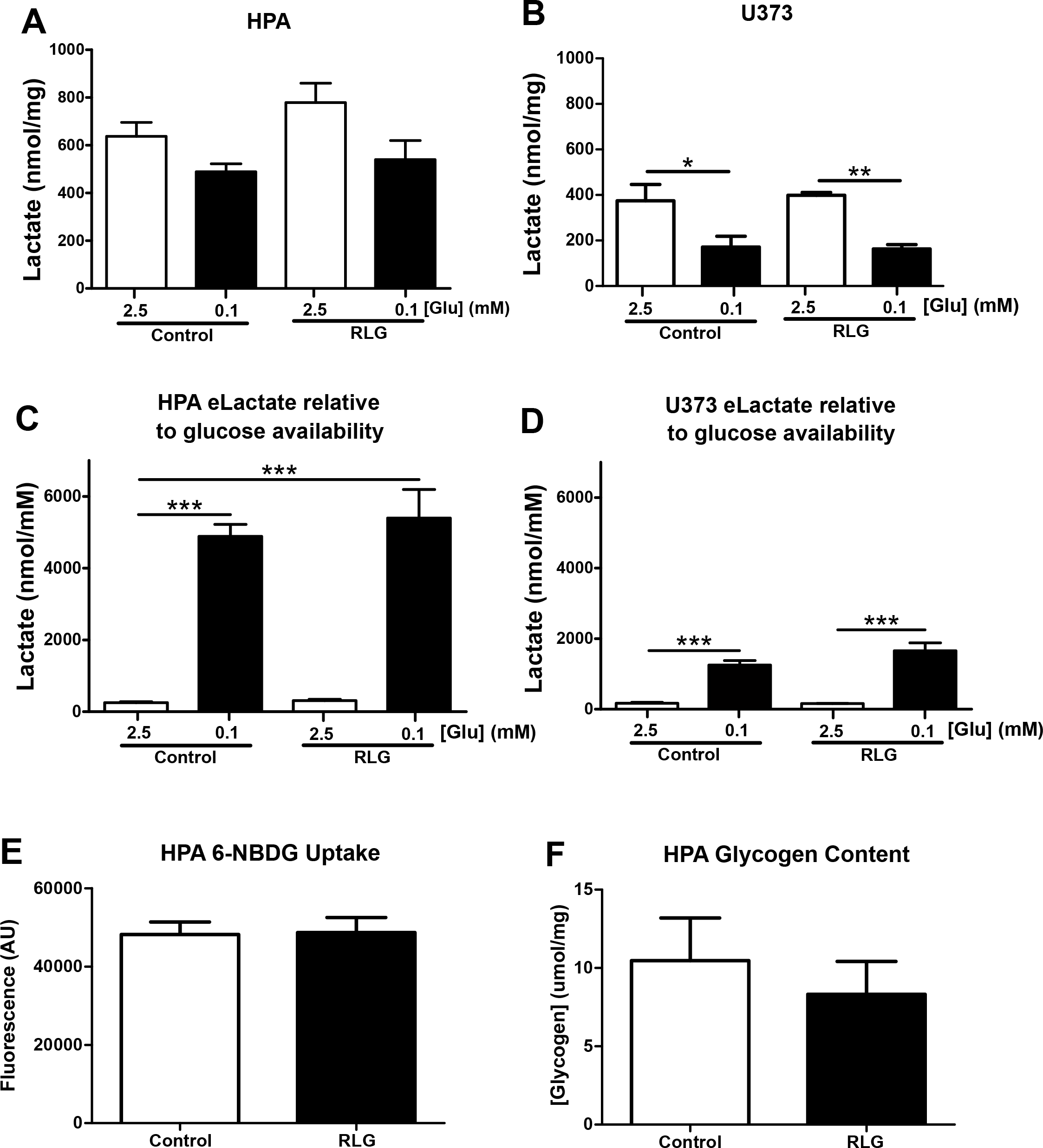
Effect of recurrent low glucose (RLG) on human astrocyte lactate release, glucose uptake and glycogen levels. Extracellular lactate levels measured from conditioned media from human primary astrocytes (HPA; **A**; n=8) and U373 (**B**; n=6) following exposure to 2.5 or 0.1 mmol/l glucose containing media for 3 hours. Extracellular lactate levels normalized to glucose availability from conditioned media from HPA (**C**; n=8) and U373 (**D**; n=6) following exposure to 2.5 or 0.1 mmol/l glucose containing media for 3 hours. ***E***. Fluorescent signal from labeled glucose analogue 6-NBDG incubated with HPA cells for 15 minutes. ***F***. Glycogen content of HPA cells in control and following exposure to RLG (n=5). \**P*<0.05; \*\**P*<0.01;\*\*\**P*<0.001.

We next assessed the degree of glucose uptake by human astrocytes using the fluorescently labelled glucose analogue, 6-(N-(7-Nitrobenz-2-oxa-1,3-diazol-4-yl)amino-6-Deoxyglucose (6-NBDG). HPA cells exposed to RLG showed no change in 6-NBDG uptake (Fig. 6*E*), indicating that the persistent changes in mitochondrial oxygen consumption were not simply mediated by increased glucose uptake. Correspondingly, we did not observe any changes in astrocytic glycogen content following RLG (Fig. 6*F*).

Correlating with our mitochondrial imaging studies, the levels of mitochondrial markers succinate dehydrogenase (SDH) and the voltage-dependent anion channel (VDAC) were not altered by RLG, suggesting no change in total mitochondrial content (Fig. S5). Fatty acid synthase (FAS) and carnitine palmitoyltransferase 1a (CPT-1a) expression were also not altered, indicating that changes in FA-dependency following RLG are likely to be mediated by metabolic flux rather than expression of enzymes within the FAO pathway.

## DISCUSSION

Our studies shed new light on the previously uncharacterized intrinsic metabolic changes in human glial cells in response to acute and recurrent low glucose. We demonstrate that astrocytes, in the absence of physical and chemical signals from neurons, react to low glucose by increasing the activation of the key metabolic sensor AMPK. Changes in AMPK pathway phosphorylation occur within a pathophysiologically-relevant glucose concentration range, similar to that which affects CNS glucose-sensing neurons [28, 29].

It is well-accepted that astrocytes are more metabolically flexible than neurons, partly due to the higher expression of PFKFB3, which can accelerate glycolysis during stress [22]. Therefore, to examine specifically the changes in astrocyte metabolism following recurrent low glucose, we measured markers of mitochondrial and glycolytic metabolism in human primary astrocyte mono-cultures. Importantly, we found that daily prior bouts of low glucose (4 × 0.1 mmol/l glucose) lasting 3 hours, were sufficient to increase basal mitochondrial and glycolytic metabolism and sustain intracellular ATP levels. This increased basal mitochondrial metabolism is likely mediated, at least in part, by an increased contribution of FAO to basal oxygen consumption. Correlating with this hypothesis, we noted increased mitochondrial proton leak and reduced mitochondrial coupling efficiency in astrocytes exposed to RLG. Importantly, FAO is known to increase uncoupling, as FAO generates increased levels of reactive oxygen species (ROS)/superoxide [31] that can increase proton leak via activation of uncoupling proteins or the ATP/ADP antiporter (ANT; [32]). Previous work has demonstrated that *in vivo*, genes for β-oxidation are induced by acute hypoglycemia but this fails following recurrent hypoglycemia (RH)[33]. Our data, in astrocyte monocultures contrasts with these measurements taken from whole rodent mediobasal hypothalamus, which include both neurons and glia. Therefore, it is possible that cell type-specific changes are masked when measuring mRNA expression from whole tissue sections. Despite the evidence of mitochondrial stress and in contrast to our expectation, examination of the filamentous mitochondrial networks revealed no change in either mitochondrial size or number following acute low glucose or following RLG. Indeed, mitochondria exposed to RLG generally displayed a normal filamentous network, with little or no evidence of mitochondrial fragmentation. These data suggest that astrocyte mitochondria successfully adapt to maintain energy production in response to repeated bouts of low glucose availability.

Previous studies in rats found no change in brain glycogen levels following RH [34] or any correlation between brain glycogen content and impaired awareness of hypoglycemia in humans [35]. However, other studies suggest brain glycogen content is increased following recovery from insulin-induced hypoglycemia [36] and following recurrent 2-DG induced glucoprivation in mice [37]. Our data clearly demonstrate that astrocytes intrinsically adapt by increasing cellular metabolism in response to recurrent energy stress. Specifically, rates of extracellular acidification (a marker for glycolysis) following glucose “reperfusion” were elevated in cells exposed to RLG, suggesting a augmented re-activation of glycolysis. However, despite glycolysis being a major determinant of glycogen synthesis, we found no significant change in astrocytic glycogen levels following RLG. In our model, we measured extracellular lactate accumulation (which can be generated from glycogen breakdown during aglycemia [18]) in conditioned media from astrocytes over a 3 hour period. After correcting for glucose availability, lactate release increased substantially during low glucose exposure, although the magnitude of this response was not significantly altered following RLG. We cannot rule out, however, the possibility that more subtle changes in lactate and glycogen occur during the transition to low glucose or on glucose recovery. Moreover, before definitively ruling out a role for astrocytic glycogen and subsequent lactate release, a more replete *in vitro* model, containing both astrocytes and neurons should be tested, thus facilitating the inclusion of neurotransmitter dynamics into the model. The ability to produce increasingly credible human neurons from iPSC sources could potentially enable such investigations to be performed in an *in vitro* human system.

Astrocytes play a key role in maintaining glutamatergic neurotransmission through the glutamate-glutamine cycle (for recent review see [38]). Importantly, astrocytic glutamate clearance requires co-transport of three Na^+^ ions [39], meaning that, via the ATP-dependent Na^+^/K^+^ ATPase, there is a significant metabolic cost for astrocytes to maintain their Na^+^ homeostasis, estimated to be approximately 20% of astrocyte ATP production [40]. This process may become particularly costly during hypoglycemia. Moreover, glutamate stimulation of astrocytes produces mitochondrial dysfunction, decreasing mitochondrial spare respiratory capacity and increasing lactate production [41]. Furthermore, recent observations suggest that astrocytic glutamate recycling is diminished following RH, a change most likely mediated by reduced glutamate uptake [42]. Our data add to these findings and suggest that metabolic changes within astrocytes, could play a part in defective counterregulatory responses following recurrent hypoglycemia. In support of this, astrocytes are increasingly recognized as playing an important role in regulation of whole body metabolism. For example, chemogenetic activation (by increasing intracellular Ca^2+^) of astrocytes in the arcuate nucleus increased feeding in mice and conversely, sequestering astrocytic Ca^2+^, decreased food intake [15]. Moreover, modulation of inflammatory signaling in astrocytes also modulates feeding in response to a high fat diet [43, 44], further supporting a role for astrocytes in regulating energy sensing neurons of the hypothalamus. Within the hindbrain, astrocytes can detect hypoglycemia to regulate adjacent neurons to increase gastric emptying in response to a glucoprivic stimulus [14]. Astrocytes have also been shown to decrease hyperglycemia in a rodent model of type 1 diabetes [45] suggesting that astrocytes may play a fundamental role in regulating whole body glucose levels. In conclusion, our data further support the evidence that glial cells are likely to play a role in detecting hypoglycemia and that specifically changes in astrocyte mitochondrial function (increased fatty acid oxidation) could play a role in defective glucose counterregulation. This study also highlights the need for more investigation into neuron-glial interactions during and following recurrent hypoglycemia.

## Acknowledgements

This study was funded by grants from Diabetes UK (RD Lawrence Fellowship to CB; 13/0004647), the Medical Research Council (MR/N012763/1) to KLJE, ADR and CB, a Mary Kinross Charitable Trust PhD studentship to CB, ADR and RW to support PWP. Additional support for this work came from awards from the British Society for Neuroendocrinology (to CB and KLJE), Society for Endocrinology (CB), Tenovus Scotland (CB) and the University of Exeter Medical School (CB and KLJE). AR was also supported by a Royal Society Industry Fellowship. We also wish to thank Nicholas Gutowski and Janet Holley for kindly gifting the human primary astrocytes. We also wish to thank Nick Howe for guidance with Extracellular Flux Analysis. Parts of this study were presented at the American Diabetes Association 77^th^ Scientific Sessions and at the Diabetes UK Annual Professional Conference 2017.

## Author Contributions

P.G.W.P, J.M.V.W., J.L.R., R.W., A.D.R. and C.B. researched data. J.K.C., A.D.R.,
P.G.W.P, K.L.J.E., and C.B. contributed to experimental design, analysis and writing of the manuscript. C.B. conceived the study and is the guarantor of this work and, as such, had full access to all the data in the study and takes responsibility for the integrity of the data and the accuracy of the data analysis.

---

**Supplementary Figure 1. Illustration of the recurrent low glucose model**. Over 4 days, cells are exposed to 0, 1, 3 or 4 bouts of 0.1 mM glucose, representing control, acute low glucose, antecedent low glucose or recurrent low glucose, respectively.

**Supplementary Figure 2. Acute and recurrent low glucose does not alter mitochondrial number or length. A**. Representative confocal images of human primary astrocytes (HPA) cells exposed to 2.5 or 0.1 mM glucose for 3 hours following control or recurrent low glucose exposure (RLG). Raw confocal images are shown on the left hand side, with extracted signal images shown on the right. **B**. Quantification of median object size (in pixels) using a custom MatLab script (15 images across 3 separate experiments).

**Supplementary Figure 3. Recurrent low glucose (RLG) does not alter glycolytic re-activation after acute low glucose exposure in U373 cells**. Extracellular acidification rates (ECAR) increased in U373 cells on re-introduction of 0.5 mmol/l glucose (**A**), 2.5 mmol/l glucose (**B**) and 5.5 mmol/l glucose (**C**) although this was not significantly different between control and recurrent low glucose (RLG) treated cells. Oxygen consumption rate (OCR) decreased on addition of 0.5 mmol/l glucose (**D**), 2.5 mmol/l glucose (**E**) and 5.5 mmol/l glucose (**F**), which was not significantly different between control and RLG (n=8-10).

**Supplementary Figure 4. Expression of key mitochondrial and glycolytic markers are not altered by acute or recurrent low glucose exposure (RLG)**. Glucose transporter 1 (GLUT-1), hexokinase I (HKI), hexokinase II (HKII), hexokinase III (HKIII), phosphofructokinase [platelet; PFK(P)], voltage dependent anion channel (VDAC), succinate dehydrogenase A (SDHA), carnitine palmitoyltransferase 1α (CPT1α) and fatty acid synthesis (FAS) were not altered by acute or recurrent low glucose exposure (n=3-4).

